# Quaternary climatic changes and biogeographic barriers drove codiversification in the obligate mutualism between *Camponotus laevigatus* and its endosymbiont *Blochmaniella*

**DOI:** 10.64898/2026.04.02.716133

**Authors:** Swapnil S. Boyane, Garrett J. Behrends, Joseph D. Manthey

## Abstract

Codiversification often arises when hosts and their endosymbionts share a linked evolutionary history, exhibit vertical transmission, or share ecological and biogeographic processes. Most studies on the codiversification of carpenter ants (genus *Camponotus*) have focused on the co-phylogeny of hosts and endosymbionts across multiple species; however, no studies have examined the intraspecific population-level phylogeographic patterns of codiversification within *Camponotus*. California has been a geographic focus for phylogeographic studies due to its high endemism and complex geographic structure, and *Camponotus laevigatus* is a carpenter ant primarily found there. Here, we used whole-genome sequencing from *C. laevigatus* and its endosymbiont, *Blochmaniella* to investigate phylogeographic patterns of host-endosymbiont codiversification and estimated kinship of ants sampled near one another. We identified three phylogeographic clusters and isolation-by-distance analyses indicated a positive relationship between genetic and geographic distance in *C*. *laevigatus* and *Blochmaniella*. Using estimates of effective migration surfaces, we found that the Central Valley in California acts as a significant barrier to gene flow among populations. Our phylogenetic analyses revealed the congruent phylogenies of *C*. *laevigatus* and *Blochmaniella*, supporting codiversification. We also estimated kinship among individuals from the same and nearby sampling sites; kinship results indicated full-sister relationships among individuals from the same sampling site, except for three pairwise comparisons, and foragers from nearby sampling sites displayed some shared kinship. Lastly, our demographic analysis revealed a Pleistocene divergence, highlighting the role of Quaternary climatic cycles in shaping the population structure of *C*. *laevigatus*.

## Introduction

Insects, one of the most speciose taxa on the planet with an evolutionary history of ∼479 million years (Misof et al., 2014), are home to a wide variety of microbes and have developed associations with microbial partners that have led to novel evolutionary innovations (Koga et al., 2013; Sudakaran et al., 2017). These interactions between microorganisms and their hosts are diverse; with some bacteria forming beneficial mutualistic associations (mutualism), while others are harmful to hosts (parasitism) (Kikuchi, 2009). Microbial symbionts often confer a range of benefits to their hosts, including increased resistance to infections, abiotic stresses, and chemical pesticides. (Heyworth & Ferrari, 2016; Kikuchi et al., 2012). Among these symbionts, obligate endosymbionts—bacteria that live within the host’s body, sometimes within specialized cells, and that are often passed on from mother to offspring—play important roles for their host by providing nutrition and helping in metabolic processes (Gil et al., 2003; Kikuchi, 2009; McCutcheon et al., 2019; Wernegreen, 2002) and protecting against pathogens (Nakabachi et al., 2013). Obligate endosymbionts have played a significant role in the evolutionary success of approximately 10% of insect species (Wernegreen, 2002) and often influence host evolution, leading to lineage codiversification (Bennett & Moran, 2015; Moeller et al., 2023). Therefore, understanding the patterns and drivers of codiversification between hosts and their obligate endosymbionts remains a central question in evolutionary biology.

Codiversification often arises when hosts and their symbionts share a linked evolutionary history, resulting from maternal inheritance, long-term symbiosis, or shared ecological and biogeographic processes (De Vienne et al., 2007; de Vienne et al., 2013; Fromont et al., 2016; Moeller et al., 2023; Moran et al., 2008; Moran & Sloan, 2015). Codiversification may be favored by selection or influenced by the limited dispersal of hosts, as new lineages adapt to new niches (Rundell & Price, 2009). Dispersal barriers play a critical role in codiversification, as they can limit gene flow and promote reproductive isolation among hosts (Sobel et al., 2010). To maintain these patterns of codiversification, both the geographic and reproductive isolation of the host and their endosymbiont lineages are important (Suzuki et al., 2022). For example, in cases of maternally inherited obligate endosymbionts, dispersal of the endosymbiont is inherently limited by host movement, which contributes to codiversification between hosts and their endosymbionts (Clayton et al., 2015). Moreover, previous studies have documented patterns of codiversification in diverse insect groups, such as Hemiptera (Fromont et al., 2016; Liang et al., 2024; Urban & Cryan, 2012), Coleoptera (Wierz et al., 2024), Hymenoptera (Cruaud et al., 2012), and Lepidoptera (Russell et al., 2012). However, incongruence can arise when endosymbionts undergo host switching (Clayton et al., 2015; Hayward et al., 2021) or exhibit deep coalescence relative to the host phylogeny. Understanding the processes that lead to congruent or incongruent host-endosymbiont phylogenies at a shallow evolutionary timescale is strengthened with a phylogeographic approach. The phylogeographic framework offers a powerful paradigm for studying codiversification by investigating how geographic isolation and historical climatic events shape the genetic structure and patterns of codiversification in hosts and their endosymbionts.

Hymenopterans, which include ants, bees, and wasps, are among the most diverse and speciose insect groups and play significant ecological roles (Grimaldi & Engel, 2005). The carpenter ant genus *Camponotus* Mayr, 1861, is the most species-rich genus in the tribe Camponotini with more than 1,500 described species spread across approximately 46 subgenera worldwide (Rossi & Feldhaar, 2020). The North American subgenus *Camponotus* (*Camponotus*) includes species commonly found in woodlands (Mackay, 2019). One such species, *Camponotus laevigatus* (Smith, F., 1858), is primarily found in California, where they usually nest in dead branches of live oak (*Quercus*) trees (Gadau et al., 1999). This genus harbors a vertically transmitted bacterial endosymbiont known as *Blochmaniella* within specialized cells called bacteriocytes, primarily located in the gaster of all castes as well as the oocytes of queens (Ramalho et al., 2018; Wernegreen et al., 2009). This intracellular, obligate endosymbiont was first discovered in the genus *Camponotus* by Friedrich Blochmann in 1887 (Blochmann, 1892). *Blochmaniella* helps their host, *Camponotus*, by providing them with essential amino acids, which play important roles in host development and nutrition throughout their life cycle (Sabree et al., 2009; Wernegreen et al., 2009). A long-term association between *Blochmaniella* and their hosts from the tribe Camponotini is estimated to span around 40 million years (Williams & Wernegreen, 2015). This mutualistic relationship has led to co-speciation, as the evolutionary history of the endosymbionts reflects that of their hosts (Degnan et al., 2004; Sauer et al., 2000; Wernegreen et al., 2009).

The Californian fauna has been the focus of phylogeographic research in recent years due to its complex geographic structure, high endemism rate, and historical geological changes. (Schierenbeck, 2014; Sork et al., 2016). For instance, during the Pleistocene epoch, around 2 million years ago (mya) to 10,000 years ago (kya), Earth’s climate experienced frequent glacial– interglacial (cold–warm) cycles, causing demographic changes in species’ populations (Hewitt, 2000; Hewitt, 2004). The most recent of these, the last glacial maximum (LGM), occurred ∼26.5 to 19 thousand years ago (Kya) (Clark et al., 2009). During this period, the global climate was at its coldest, with a large ice sheet covering much of North America, and global sea levels were at their lowest (Lambeck et al., 2014; Shafer et al., 2010). These climatic shifts played a significant role in shaping current species distributions and population structure (Hewitt, 2003).

Comparative phylogeography provides a framework for studying co-distributed taxa and addressing questions related to vicariance. Previous studies focusing on taxa found in California have identified largely congruent phylogeographic patterns and genetic differentiation associated with comparable geographic barriers, such as the Central Valley, the Sierra Nevada, and the Mojave Desert (Feldman & Spicer, 2006). For example, vertebrate taxa like snakes (Myers et al., 2013), amphibians (Kuchta & Tan, 2006), and turtles (Spinks & Shaffer, 2005) share a prominent phylogeographic break across these barriers, implying that the present-day geographic structure is mainly attributed to Pleistocene climatic cycles and past geological events. On the other hand, despite their diversity and ecological importance, the phylogeography of insect taxa in California remains underexplored. A very few studies are available, such as on beetles (Chatzimanolis & Caterino, 2007; Chatzimanolis & Caterino, 2008), moths (Rubinoff et al., 2021), ice crawlers (Schoville & Roderick, 2010), and spiders (Starrett et al., 2024). To date, no studies in the genus *Camponotus* have explored within-species phylogeographic patterns of codiversification between the host and its endosymbiont. The mutualistic association between *C. laevigatus* and its endosymbiont *Blochmaniella* and their geographic distribution across fragmented oak forests in California make this system ideal for investigating phylogeographic patterns of codiversification between hosts and their endosymbionts.

Here, we used whole-genome sequencing data from 29 individuals spanning the entire geographic range of *C. laevigatus* to investigate phylogeographic patterns of codiversification in hosts and their endosymbionts. Our goals were to use high-resolution genomic data to examine genomic variation in both *C. laevigatus* and its endosymbiont, *Blochmanniella,* and to assess kinship among *C. laevigatus samples*. Specifically, we asked, i) does *C. laevigatus* display evidence of population genetic structure in California? ii) do the host and endosymbiont exhibit congruent phylogenies, indicating a pattern of codiversification? iii) is there gene flow among populations, and do geographical barriers, such as the Central Valley, contribute to population differentiation? iv) do individuals foraging together or nearby one another show evidence of being full sisters originating from a singly mated queen?

## Materials and Methods

### Sampling, DNA extraction, and genome sequencing

We sampled 29 individuals while they were foraging at 21 sites spanning the geographic range of *Camponotus laevigatus* (Figure 1A). Here, sampling sites were areas where multiple individuals were found foraging on the same oak tree within a 1-2 m^2^ area. Because colonies are likely found high up in live oak trees (Gadau et al., 1999), we were precluded from finding colonies in a timely and safe manner and limited our sampling to foraging individuals. We sampled one to three workers per sampling site and stored them in cryotubes in liquid nitrogen until they were transferred to a −80°C freezer for long-term storage. Additional individuals from each sampling site were sampled for pinned voucher specimens deposited in the Museum of Texas Tech University Invertebrate Zoology Collection (Table S1).

**Figure 1.**
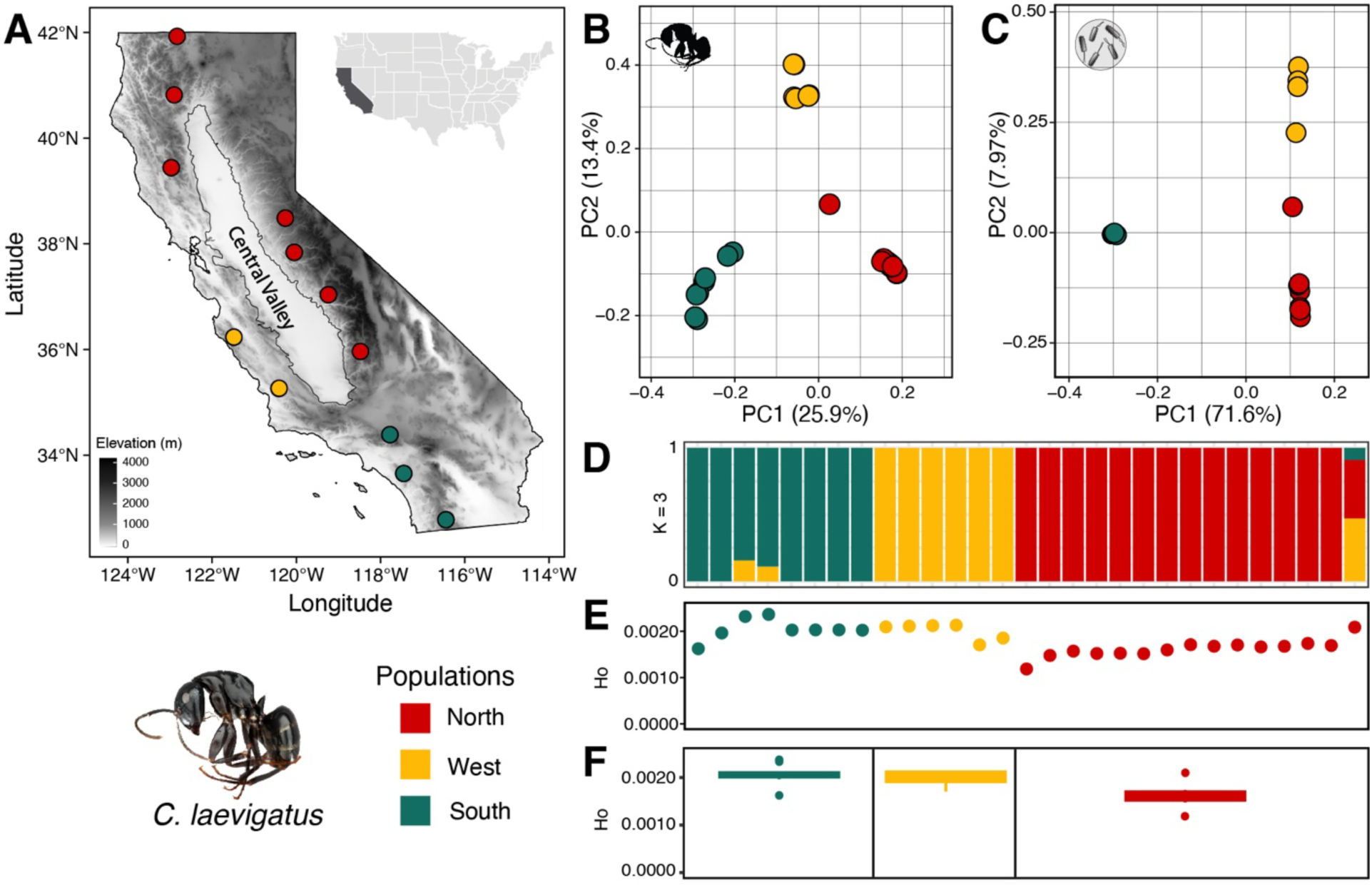
Sampling sites and geographic structure of *C. laevigatus* and its endosymbiont *Blochmaniella* across its range in California. (A) Sampling locations of 29 ants from 21 sampling sites. Different colors indicate different populations in this study (B) Principal component analysis (PCA) of the host, *C. laevigatus,* showing three genetic clusters. (C) PCA of *Blochmaniella,* which is congruent with the host PCA analysis. (D) Host population structure inferred using ADMIXTURE, considering three ancestral populations (K=3). Here, each bar represents an individual and their ancestry proportions on the Y-axis. (E) Observed heterozygosity *C. laevigatus* individuals. (F) A box plot showing the level of genetic diversity in the three populations.

To minimize excessive sequencing reads from the bacterial endosymbiont, we performed two separate extractions for each ant: (a) the head and thorax for largely host genomic DNA, and (b) the gaster for endosymbiont DNA. First, samples were pulverized in liquid nitrogen, then DNA was extracted using the Qiagen DNeasy Blood and Tissue Kit according to the manufacturer’s protocol. DNA quantification was then performed using an Invitrogen Qubit fluorometer. For each individual, the extracted DNA was pooled in a 70:30 ratio, with 70% from the head and thorax and 30% from the gaster, a protocol that has worked for our lab in past sequencing of this genus and its endosymbiont (Manthey et al., 2022). Finally, the DNA extractions were sent to the Center for Biotechnology and Genomics at Texas Tech University (Lubbock, Texas), where we outsourced library preparation and whole-genome shotgun sequencing using the Illumina NovaSeq 6000.

### Genome alignment, genotyping, and filtering *C. laevigatus*

We used the bbduk.sh script of BBMap to remove adapters and quality filter raw sequences (Bushnell, 2014). We then used the BWA-MEM algorithm of BWA v0.7.17 (Li & Durbin, 2009) to align filtered reads to a *Camponotus* sp. reference genome (Manthey et al., 2022). Next, we used Samtools v1.4.1 (Li & Durbin, 2009) to convert the SAM file to BAM format and measure sequencing coverage per individual. Then, we used the Genome Analysis Toolkit (GATK) v4.1.0.0 (McKenna et al., 2010) to clean, sort, and add read groups to the BAM files. Genotyping of these samples was done using GATK in three steps: (a) initially, we used the ‘HaplotypeCaller’ function to genotype each individual, (b) the ‘GenomicsDBImport’ function to create a database for each chromosomal segment, and (c) the ‘GenotypeGVCFs’ function to group genotypes of all individuals. Finally, we filtered genotyped sites in the host *C*. *laevigatus* using VCFtools v0.1.14 (Danecek et al., 2011) with the following flags : (a) minimum site quality of 20, (b) maximum depth (coverage) of 50, (c) minimum depth (coverage) of 6, and (e) removal of indels.

#### Blochmaniella

We first extracted putative *Blochmaniella* reads from the raw FASTQ files using the bbsplit.sh script from BBMap (Bushnell, 2014). We then separated the interleaved file using the Seqtk toolkit (https://github.com/lh3/seqtk). Next, we applied the same alignment pipeline used for the host genome to the *Blochmaniella* sequences. For genotyping, we specified “--ploidy 1” in the HaplotypeCaller and GenotypeGVCFs functions of GATK to account for the haploid nature of the *Blochmaniella* genome. Due to the low coverage (<2x) of *Blochmaniella* reads in sample C-289, we removed it from all endosymbiont analyses (Table S1).

Finally, for *Blochmaniella*, we filtered for (a) a minimum site quality of 20 and (b) a minimum depth (coverage) of 6. For downstream analyses, we used various parameters (e.g., thresholds for missing data and the inclusion/exclusion of the outgroup), as listed in Table 1.

**Table 1.**
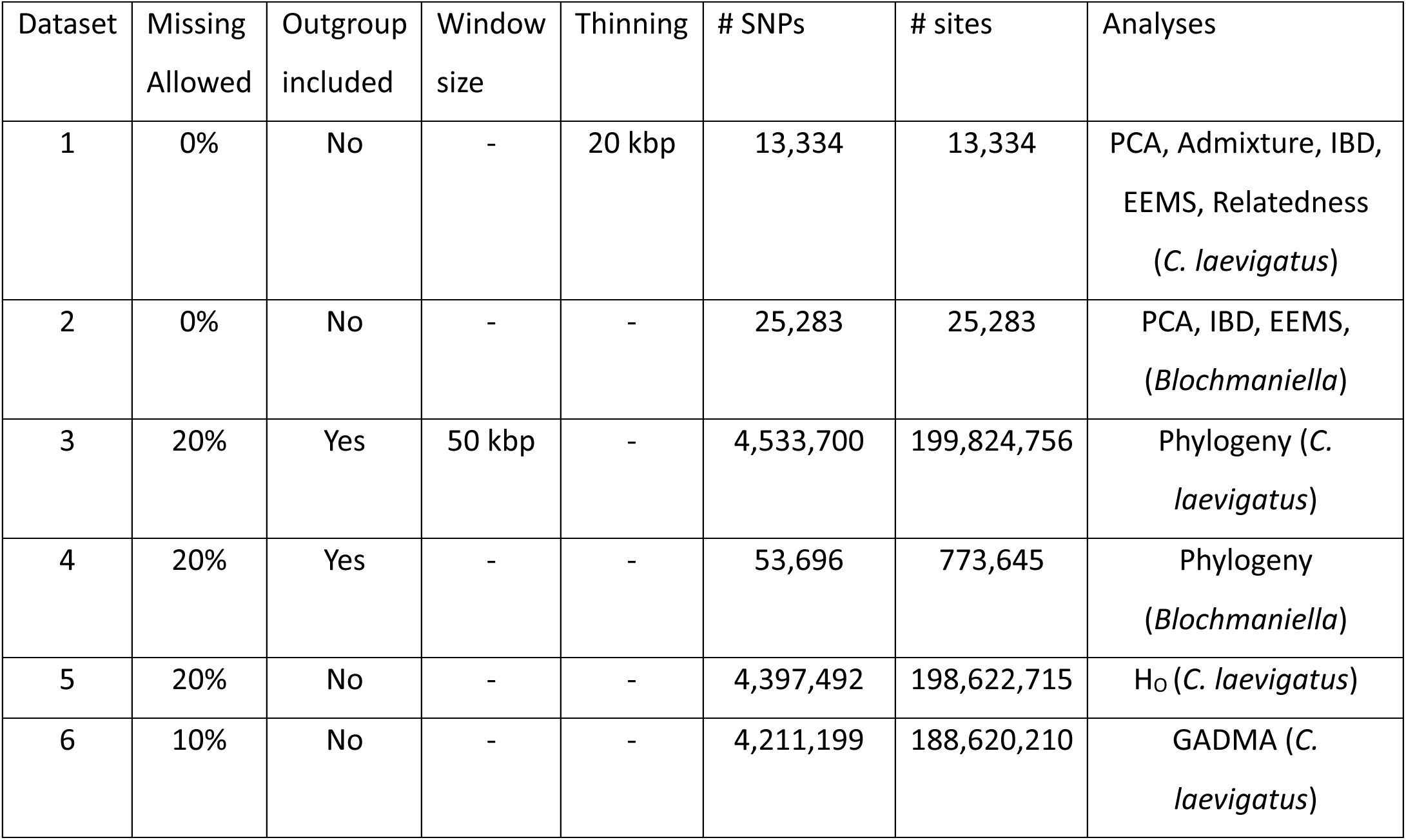
Different datasets used in this study and their characteristics.

### Population structure analyses

To estimate population genetic structure and genomic ancestry proportions for all *C. laevigatus* individuals, we used ADMIXTURE v1.3.0 (Alexander et al., 2009) with one to five genetic groups (i.e., K = 1 to K = 5). Additionally, using PLINK v2.0 (Purcell et al., 2007), we performed principal component analysis (PCA) to summarize phylogeographic structure in both *C. laevigatus* and *Blochmaniella*.

### *C. laevigatus* genetic diversity

We used observed heterozygosity (H_O_) as a measure of genetic diversity. For each individual, we measured H_O_ by dividing the number of heterozygous sites by the total number of genotyped sites passing filters.

### Phylogenomics

To infer phylogenetic relationships among the *C. laevigatus* host individuals, we estimated gene trees for 50-kbp non-overlapping windows. We further retained only windows containing at least 10,000 genotyped sites after filtering. Later, these windows were used to estimate each 50-kbp gene tree using the GTRGAMMA model of sequence evolution in RAxML v8.2.12 (Stamatakis, 2014). This generated 5324 gene trees, which were later summarized into a maximum clade credibility tree using the sumtrees.py script, a part of the DendroPy Python package (Sukumaran & Holder, 2010). Finally, we used the RaxML gene trees as input for ASTRAL III v 5.7.1 (Zhang et al., 2018) to generate a species tree. For *Blochmaniella*, we used genome-wide SNP data and the Python script vcf2phylip v2.0 (Ortiz, 2019) to convert VCF-formatted SNPs to PHYLIP format. Then, we used RAxML v8.2.12 (Stamatakis, 2014) under the GTRGAMMA model of sequence evolution to estimate the phylogenetic tree. Finally, we used FigTree v1.4.4 (Rambaut, 2007) to visualize and root the phylogenetic trees.

### Isolation by distance (IBD)

We measured the correlation between genetic and geographic distances for all pairwise comparisons among individuals to test for isolation-by-distance (IBD). We assessed whether there was a significant signal of isolation-by-distance (IBD) within the entire species and within each population cluster identified by ADMIXTURE. For both *C. laevigatus* and *Blochmaniella,* we calculated pairwise genetic distances using the “StAMPP” package (Pembleton et al., 2013), while pairwise geographic distances were computed using the “fields” package (Nychka et al., 2015) in R v4.4.0 (R Core Team, 2023). Finally, to investigate the relationship between genetic and geographic distances, we performed a Mantel test. This statistical method evaluates the correlation between two distance matrices, using the “ade4” package (Dray & Dufour, 2007) with 10,000 permutations.

### Estimation of effective migration surfaces (EEMS)

To infer population genomic connectivity across the landscape, we used Estimating Effective Migration Surfaces (EEMS) (Petkova et al., 2016). EEMS highlights potential corridors or barriers to gene flow by modeling migration rates among demes (Petkova et al., 2016). We used a similar workflow for both *C. laevigatus* and *Blochmaniella*, except for one adjustment for *Blochmaniella* in the ploidy assignment, where we specified “diploid = false.”

Next, we used the bed2diffs pipeline in R v4.4.0 (R Core Team, 2023) from the EEMS package to compute pairwise genetic dissimilarities and generate the genetic distance matrix. The analysis used the default number of MCMC iterations (2,000,000) and 1,000,000 burn-in iterations, with 200 demes and a thinning interval of 9999. We verified for MCMC convergence by visualizing the resulting MCMC trace. Finally, we visualized the results using the rEEMSplots (https://github.com/dipetkov/reemsplots2) package in R v4.4.0 (R Core Team, 2023).

### Demographic inference

To estimate divergence times among the identified phylogeographic lineages and infer population size changes over time, we used GADMA version 2.0.0 (Noskova et al., 2020), which employs the “moments” engine (Jouganous et al., 2017). GADMA reconstructs demographic histories for up to three populations from the joint site-frequency spectrum (SFS). First, we simulated the SFS with the “--preview” flag in easySFS (https://github.com/isaacovercast/easySFS) to determine the optimal projection values. Then, we downsampled the projection size and selected the allelic sample size that maximized the number of segregating sites. Individuals with admixed ancestry were excluded from the analysis. For calculating absolute divergence times, GADMA requires the mutation rate. We used a mutation rate of 1.9×10^-9^ based on a previous study (Manthey et al., 2022). The total sequence length included both variant and invariant sites. Because the generation time in this system is not known and may vary based on the age of reproductive maturity, we explored a range of generation times (5–7 years) to assess the divergence-time estimates. We ran 10 repetitions and selected the best-fit model using the highest log-likelihood value.

### Relatedness and kinship

To infer kinship among individuals, we used an allele-frequency-free method based on three ratios: R0, R1, and the KING-robust kinship estimator (Manichaikul et al., 2010; Waples et al., 2019). Each of these ratios represents a different pattern of genome-wide allele sharing between two diploid individuals (Waples et al., 2019). R0 measures the proportion of sites where both individuals have different homozygous genotypes (i.e., fixed differences), while the R1 ratio measures the proportion of sites where both individuals are heterozygous. The KING-robust kinship estimator uses allelic sharing combinations that partially overlap with those used in R0 and R1 to provide a continuous measure of kinship in the presence of population structure (Waples et al., 2019).

We categorized individuals based on their kinship metrics: pairs with a KING-robust kinship > 0.3 and an R0 ratio < 0 were classified as possible full sisters (FS). Furthermore, pairs with a KING-robust kinship of 0.15 to 0.3 were classified as half-siblings (HS), pairs with a KING-robust kinship of 0.10 to 0.14 were classified as distantly related (DR), and pairs with a KING-robust kinship of less than 0.10 were classified as unrelated (UR). Next, we grouped samples into three types of pairwise comparisons to evaluate kinship: (1) foragers from same sampling site, which involved individuals from the same sampling location (i.e., foraging together on a tree trunk), (2) foragers from nearby sampling sites, using individuals from sampling locations situated within 1 km of each other, and (3) foragers from distant sampling sites, involving individuals located more than 1 km apart (Figure 4A). Finally, we created a heatmap using the R packages “pheatmap” (https://github.com/raivokolde/pheatmap) and “tidyverse” (Wickham et al., 2019).

## Results

### Genetic structure and diversity of the host (*C. laevigatus*)

The PCA of *C. laevigatus* showed three genomic clusters corresponding to the geographic distribution of the samples (Figure 1B). These clusters are hereafter referred to as the northern, western, and southern populations. PC1 separated all three genetic clusters, while PC2 separated the western population from the southern and northern populations (Figure 1B). We observed a close clustering within the populations, except for the northern population, where one outlier was positioned intermediately between the clusters of the other regions.

ADMIXTURE analysis aligned with the genetic structure found in the PCA analysis. The optimal value of K for the dataset was determined to be K = 3, based on the PCA results, known geographic distribution, and cross-validation scores. The three genetic clusters again corresponded to the samples’ geographical distribution. Additionally, three individuals were found as potentially admixed—two from the southern populations and one from the northern population (Figure 1D).

Consistent with PCA and ADMIXTURE results, both the consensus RAxML maximum-likelihood tree and the ASTRAL-III tree recovered three major clades corresponding to geographic regions (Figure 3). Individuals from the northern populations formed a distinct clade, with an admixed individual (C-294) from this group forming an independent lineage. The northern population was sister to the western population, and together, they were sister to the southern population (Figure 3).

Our overall dataset revealed a significant pattern of isolation by distance (IBD), indicating that genetic differentiation increases with geographic distance (r = 0.7331, p < 0.001).

Furthermore, each regional group identified by ADMIXTURE showed a positive relationship between genetic and geographic distances (Figure 4A). The southern populations showed the strongest IBD pattern (*r* = 0.869, *p* = 0.002), followed by the western (*r* = 0.682, *p* = 0.023) and northern (*r* = 0.516, *p* < 0.001) populations. Similarly, EEMS analysis provided a spatial view of gene flow, indicating a corridor for higher gene flow among northern sampling sites; however, relatively lower gene flow in the southern and western populations (Figure 4C). This analysis suggested that the Central Valley is a significant barrier to gene flow between *C. laevigatus* populations.

### Genetic diversity and demographic histories of the host

H_O_ varied across the three populations; the southern population showed the highest H_O_, ranging from 0.0016 to 0.0023, with two outlier individuals. The western population had H_O_ ranging from 0.0017 to 0.0021 (Figure 1E and 1F). The northern population showed the lowest H_O_, ranging from 0.0011 to 0.0017, with one outlier around 0.0021. All outliers were identified as potential admixed individuals in genetic structure analyses. To further explore geographic patterns, we plotted H_O_ vs. latitude, which showed a negative relationship: as latitude increased, H_O_ decreased (Figure 2).

**Figure 2.**
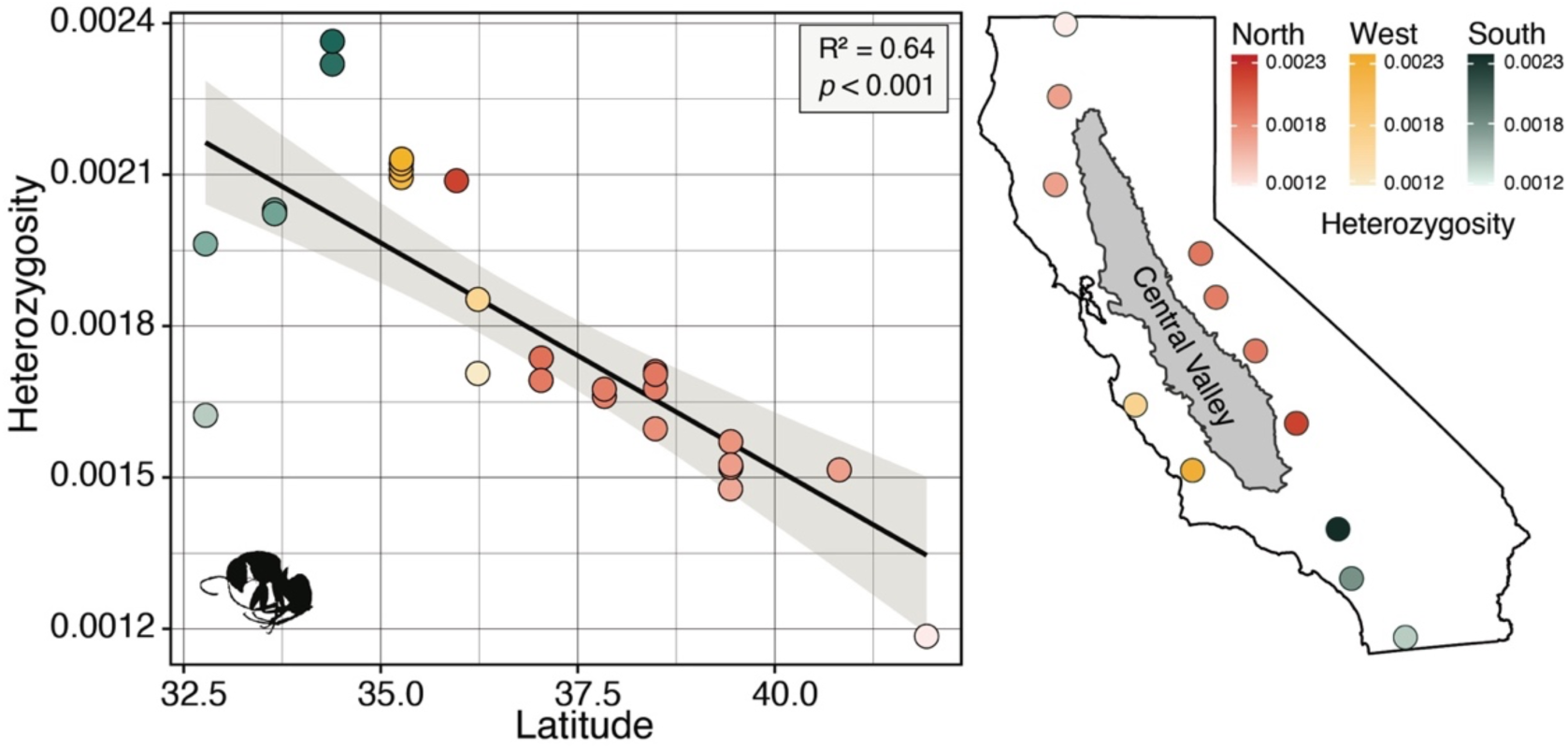
Genetic diversity and latitude relationship. Correlation between observed heterozygosity and latitude (left). Heatmap showing spatial variation in H_O_ across California (right)

**Figure 3.**
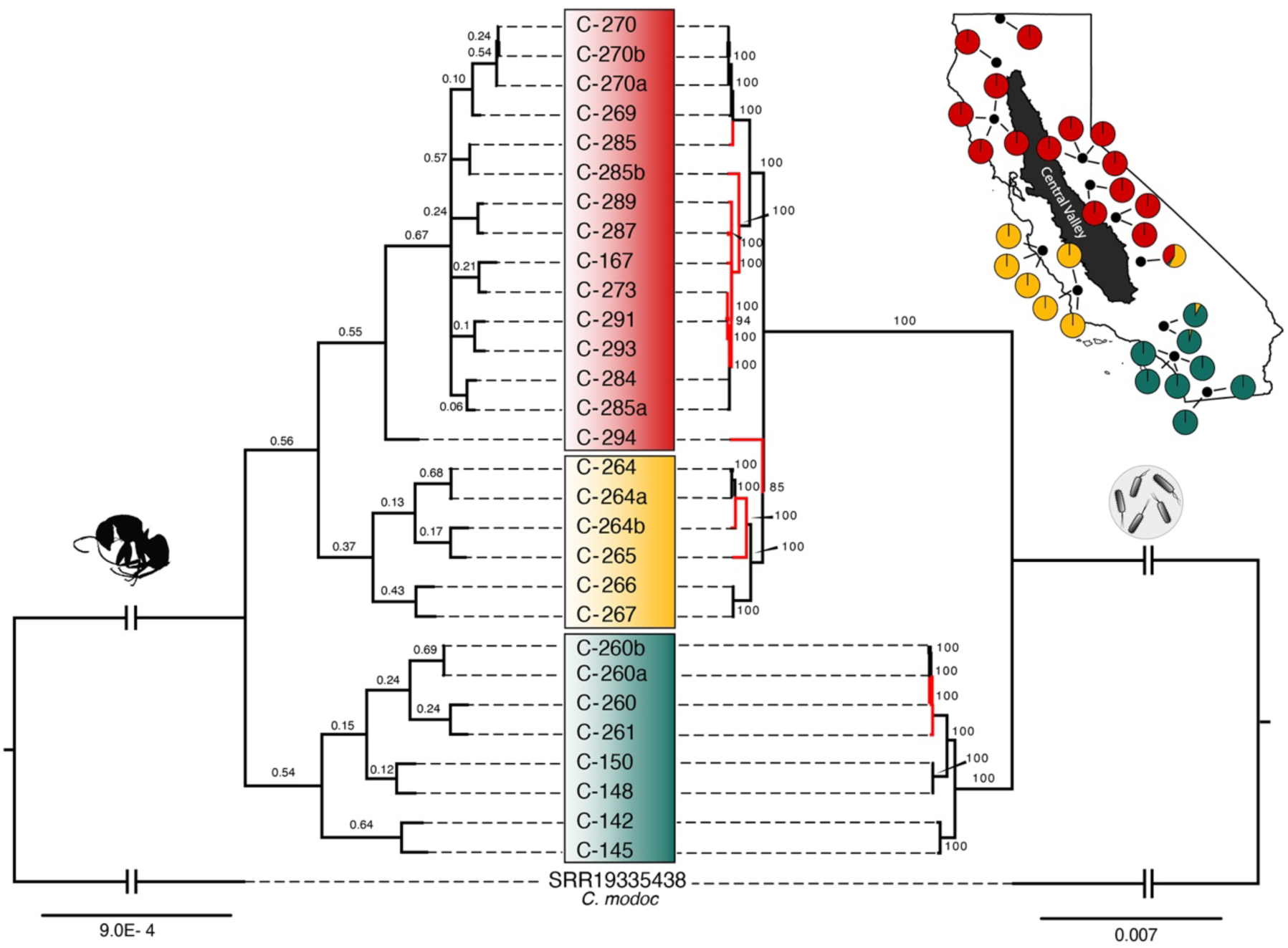
Phylogenetic congruence between *C. laevigatus* (left) and its primary endosymbiont, *Blochmaniella* (right). Both trees were rooted using the *Camponotus modoc* sample. Red-colored branches indicate discordance between the host and endosymbiont phylogenies. The map in the right corner displays sampling localities that show results of ADMIXTURE analysis with K = 3. Note: sample C-289 was excluded from the *Blochmaniella* study due to low coverage.

**Figure 4.**
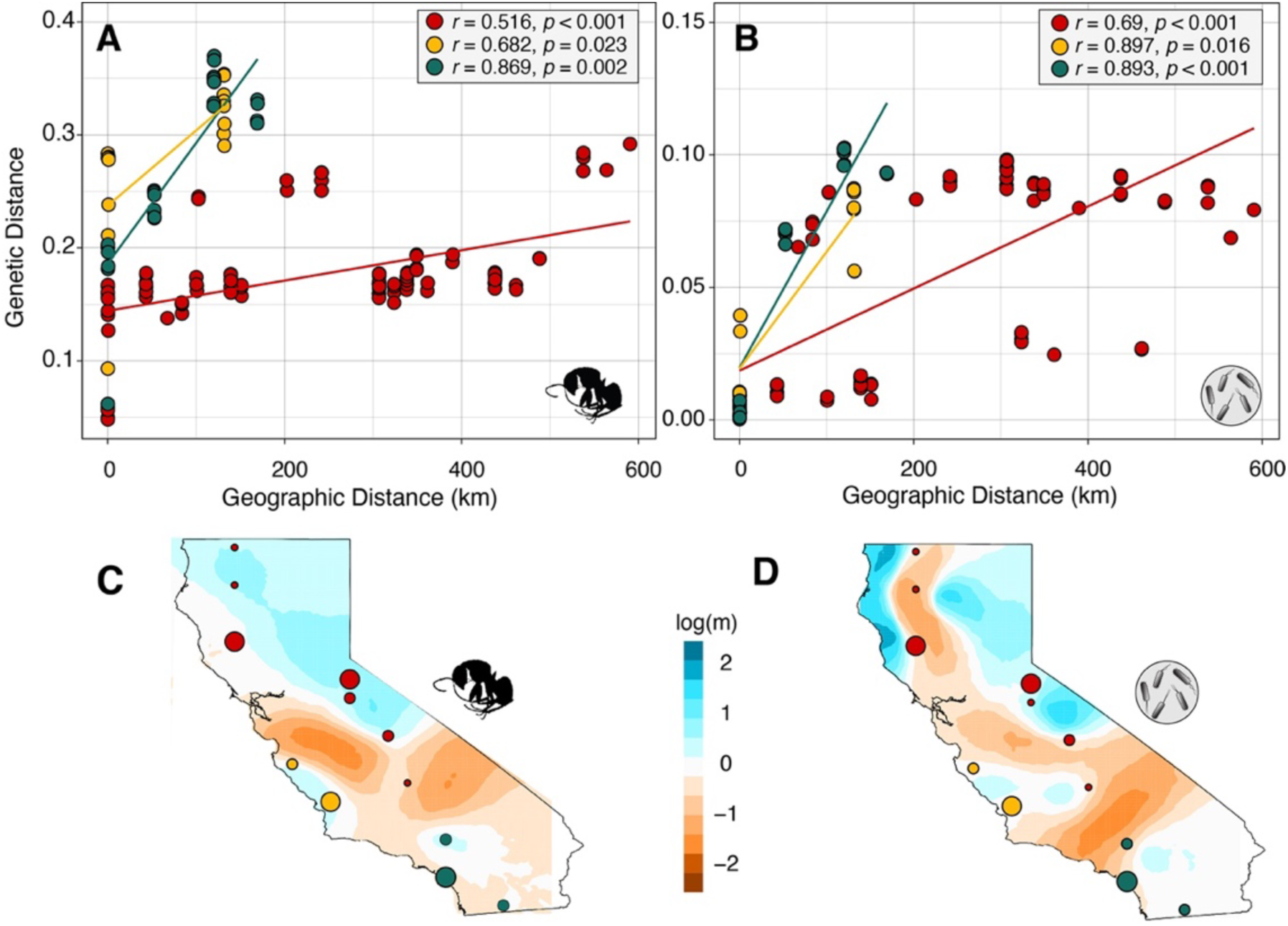
Isolation-by-distance (IBD) and estimates of effective migration surfaces (EEMS) of the host and the endosymbiont. (A) Correlation between genetic distance and geographic distance of the host, *C*. *laevigatus*, across three populations. (B) IBD pattern of *Blochmaniella* for the same three populations. (C + D) EEMS analysis indicating effective migration surface of gene flow in (C) the host, *C*. *laevigatus*, and (D) the endosymbiont *Blochmaniella*. In these analyses, the color scale indicates gene flow: darker blue shades correspond to higher values, whereas darker red shades correspond to lower values. Note: Red, yellow, and green colors represent north, west, and south populations, respectively.

The best-fit demographic model inferred using GADMA is shown in Figure 6. It recovered two major splits: first, between the southern population and the ancestors of the northern and western populations, and a more recent divergence between the northern and western populations (Figure 6). Our model also suggested that these divergences occurred at ∼311,471 and ∼108,590 generations, respectively. After the second split, all lineages showed recent population expansion, except the Western lineage, which seems to have reduced. Our best-fit model suggested that the initial split occurred 1.5–2.1 mya, whereas a recent split between northern and western populations occurred 542–760 kya.

### Genomic structure of *Blochmaniella* endosymbionts

In the case of *Blochmaniella*, PCA showed a clustering pattern consistent with that observed in its host, in which samples from each region formed clusters, except for the northern population, where one sample was positioned apart from the others (Figure 1C).

The *Blochmaniella* gene tree, estimated using a genome-wide SNP dataset, showed a topology nearly identical to that of *C. laevigatus*, with three distinct clades. This topology was largely congruent, except for a few relationships that exhibited discordance and are highlighted in red. The majority of *Blochmaniella* nodes had 100% support (Figure 3). Similarly, our overall dataset for *Blochmaniella* indicated a positive correlation between geographic and genetic distance, as seen in *C. laevigatus* (*r* = 0.61, *p* < 0.001), indicating that genetic distance increases with geographic distance, supporting the isolation by distance hypothesis (Figure 4B). Among the three regions, the IBD pattern was strongest in the western population (*r* = 0.897, *p* = 0.016), followed by the southern population (*r* = 0.893, *p* < 0.001). The northern population showed moderate IBD (*r* = 0.69, *p* < 0.001) and additional substructure.

Finally, our EEMS analysis revealed differences in effective migration rates between endosymbionts and the host. Although the overall pattern was similar to that of the host, *Blochmaniella* showed a more pronounced barrier to gene flow, especially extending into the Klamath Mountains (Figure 4D). This more substantial or more widespread barrier to gene flow may reflect the biology of *Blochmaniella*, which is maternally transmitted, limiting its dispersal to the movement of queen hosts.

### Relatedness and kinship

Using genome-wide SNPs, we estimated kinship. To identify full-sister relationships, we expected the KING-robust kinship estimator to be > 0.3 and R0 to be < 0. As expected, six pairs of foragers collected from the same oak tree (i.e., C-260a–C-260b, C-264–C-264a, C-270–C-270a, C-270–C-270b, C-270a–C-270b, and C-285–C-285b) showed KING-robust kinship estimates ranging from 0.361 to 0.390 and R0 values between 0.000 and 0.002, consistent with full-sister relationships. Conversely, samples C-260, C-264b, and C-285a were identified as half-siblings to the other foragers found there (Figure 5C and 5D). For half-siblings, we expected KING-robust kinship values to be > 0.15 to 0.30. Among 16 pairwise comparisons of foragers from nearby trees (within 1 km), two pairs (C-142–C145 and C-266–C-267) showed KING-robust kinship values of 0.07 and 0.10, respectively, indicating somewhat related relationships (Figures 5C and 5D). Finally, we expected KING-robust kinship values to be < 0.1 for weakly related or unrelated individuals. As expected, 271 out of 351 geographically distant individuals’ pairwise comparisons showed low kinship values, suggesting that they are largely unrelated. However, 62 comparisons were identified as half-siblings, with a KING-robust kinship value ranging from 0.15 to 0.20. Among these comparisons, we observed clusters based on spatial proximity. The maximum distance between half-sibling individuals was approximately 74-276 km. The remaining 18 pairwise comparisons were distantly related, with many individuals separated by about 90 km.

**Figure 5.**
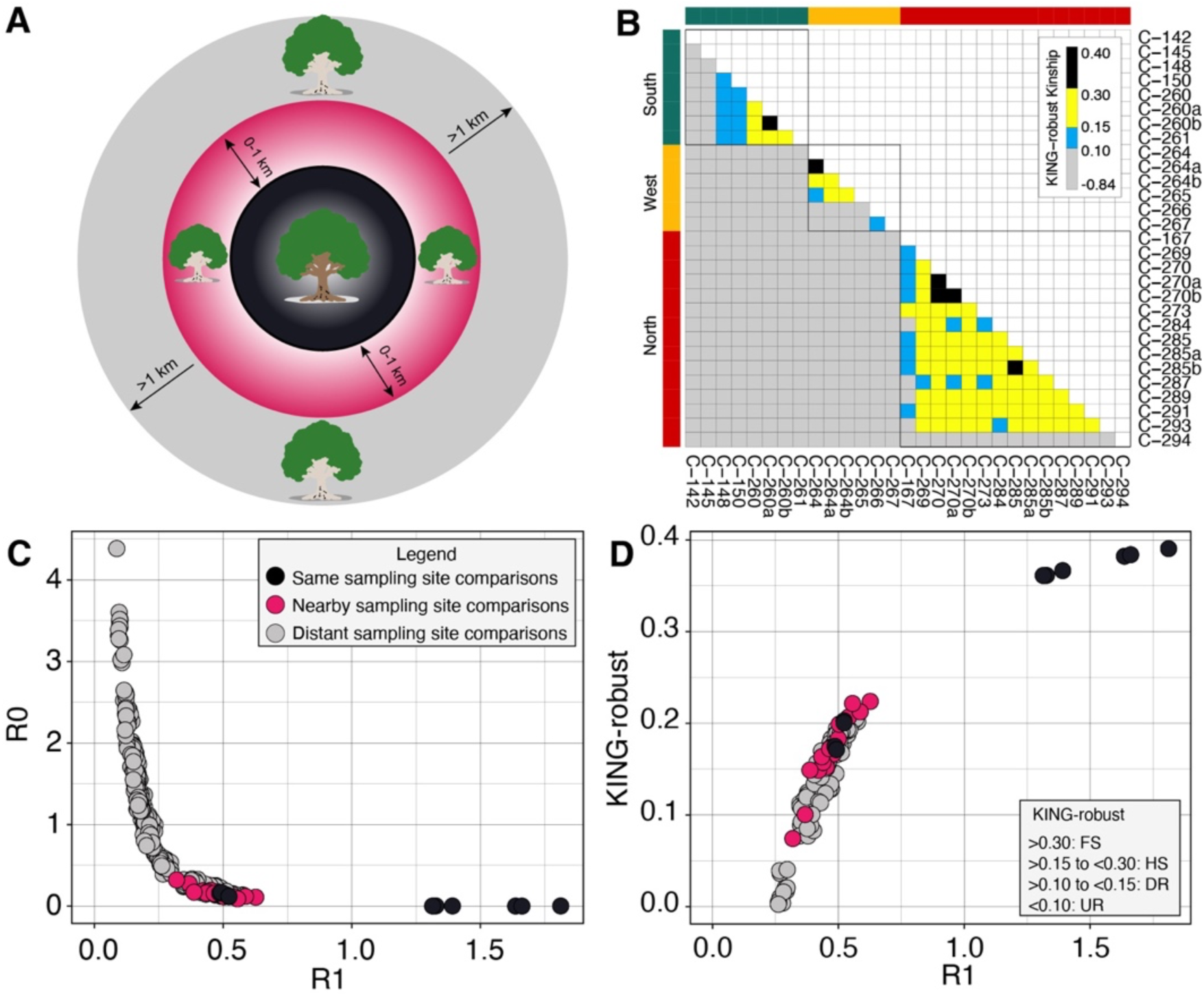
Inference of kinship based on estimates of KING-R1 and R0-R1 in *C*. *laevigatus*. (A) Graphical illustration demonstrating three levels of spatial comparisons: both individuals from the same sampling site in ∼1-2 m^2^ on a single tree trunk, colored black; comparisons where individuals were sampled at different sites but ≤1 km of each other, colored pink; and distant comparisons, which include comparisons of individuals from sampling sites that are more than 1 km apart. (B) Pairwise comparisons of KING-robust kinship estimates. (C) Relationship of R0 and R1. (D) Relationship of KING-robust kinship and R1.

**Figure 6.**
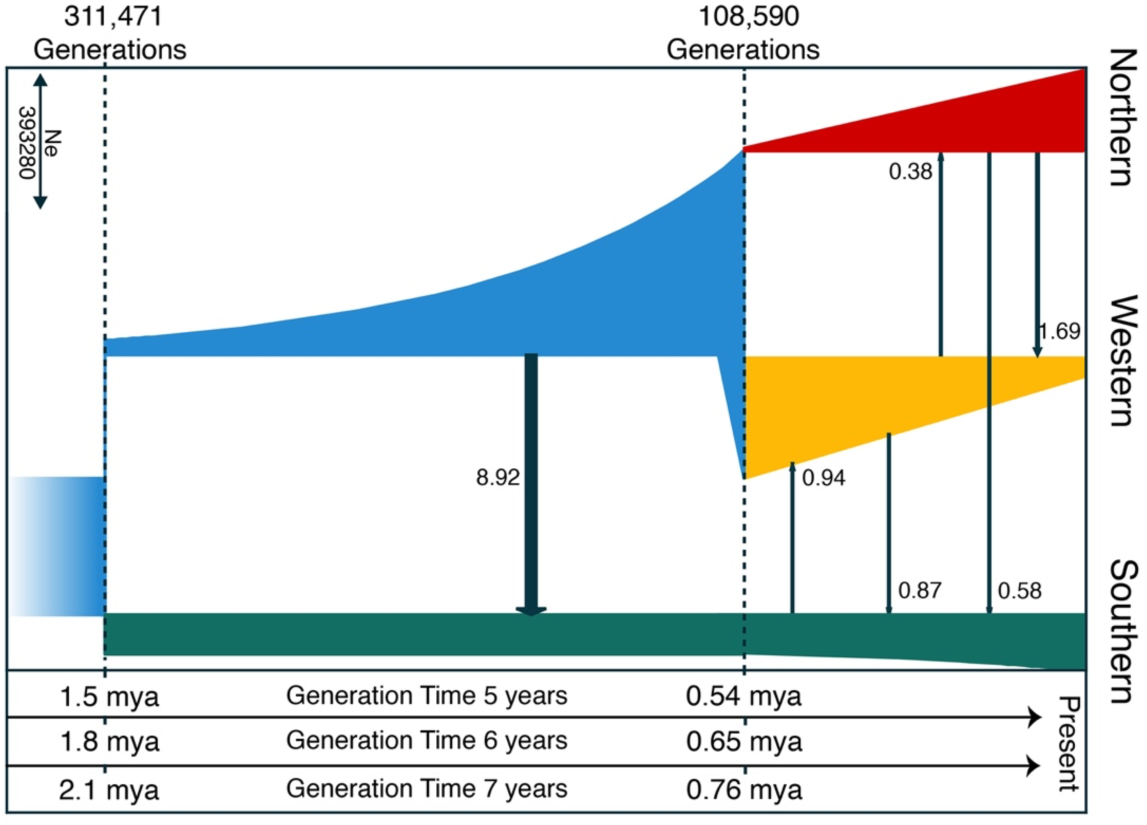
The best-fit demographic model inferred in GADMA for *C*. *laevigatus*. The top panel of the figure represents time in generations. The x-axis represents divergence time estimates (T_5_, T_6_, and T_7_) converted using generation times of 5, 6, and 7 years, respectively. Only migration edges with values > 0.1 are displayed. The units for migration edges are migrants per generation.

## Discussion

We used whole-genome sequencing data from 29 *C*. *laevigatus* individuals and their endosymbiont, *Blochmanniella*, to investigate codiversification phylogeography. Our study provides the first comprehensive insights into the phylogeographic structure of *C*. *laevigatus* and its *Blochmaniella* endosymbiont across California. By integrating analyses of population structure, gene flow, and host-endosymbiont phylogenies, we found strong but imperfect genetic clustering between the host and its endosymbiont, shaped by historical biogeographic barriers and limited gene flow.

### Influence of historical biogeographic barriers on genetic structure and gene flow in host and endosymbiont

California serves as a natural laboratory for studying phylogeography due to its complex topography, rich biodiversity, and past climatic history (Shafer et al., 2010; Sork et al., 2016). During the Pleistocene, repeated glacial oscillations caused extensive cooling and the formation of large ice sheets, forcing many species to retreat to refugia, ultimately leading to isolation among populations of different refugial regions (Batchelor et al., 2019; Hewitt, 2000; Shafer et al., 2010; Stewart et al., 2010). For example, the contraction and expansion of species’ ranges during glacial-interglacial cycles have been observed in small rodents like *Peromyscus maniculatus* (Boria & Blois, 2023). However, some wingless insects, including the ice-crawler *Grylloblatta* (Schoville & Roderick, 2010) and the ground beetle *Nebria* (Schoville et al., 2012), as well as plants like the valley oak (*Quercus lobata*) (Gugger et al., 2013), responded to these cycles by shifting from low-elevation regions to higher elevations. Such climatic oscillations, along with past geological processes, created biogeographic barriers to forest-associated taxa, including the Central Valley and the Mojave Desert, leading to shallow genetic divergence, strong population structure, and lineage diversification across several taxa (Calsbeek et al., 2003; Hewitt, 2004).

In our population structure analyses, both *C*. *laevigatus* and *Blochmaniella* indicated the presence of three phylogeographic clusters: i) a northern cluster comprising samples from the Central Sierra Nevada, north Coastal Ranges and Klamath Mountains, ii) a western cluster including samples from the Southern Coastal Ranges, and iii) a southern cluster represented by samples from the Peninsular and Transverse Ranges of California (Figure 1). Our results indicate that the observed phylogeographic structure in *C*. *laevigatus* is mainly shaped by the Central Valley, a barrier also reported in several other taxa with distributions similar to that of *C. laevigatus* but with different life histories. For example, the Central Valley is a phylogeographic barrier to plants like white oaks (Gugger et al., 2013) and the lily (Hernández et al., 2022), as well as mammals such as the dusky-footed woodrat (Boria et al., 2021), reptiles like the rubber boa (Calsbeek et al., 2003), the mountain kingsnake (Rissler et al., 2006) and the western pond turtle (Spinks & Shaffer, 2005), and birds including the wrentit (Burns & Barhoum, 2006) and the California Thrasher (Sgariglia & Burns, 2003). Furthermore, tectonic activity, such as the uplift of the coastal ranges and formation of the Transverse Ranges during the late Neogene, contributed to the Central Valley’s desiccation (Schierenbeck, 2014), creating physical barriers to migration for terrestrial taxa like reptiles and amphibians (Martinez-Solano et al., 2007; Reilly et al., 2015; Rissler et al., 2006).

In southern California, populations in the Transverse and Peninsular Ranges have shown genetic structure across several taxa, including the rove beetles (*Sepedophilus castaneus*) (Chatzimanolis & Caterino, 2007), California mountain kingsnakes (*Lampropeltis zonata*) (Myers et al., 2013; Rodríguez-Robles et al., 1999), Ring-necked snakes (*Diadophis punctatus*) (Fontanella et al., 2008), and the California newt (*Taricha torosa*) (Kuchta & Tan, 2006). On the other hand, our analyses of *C*. *laevigatus* did not show a concordant genetic break in this region, suggesting that the Transverse–Peninsular Ranges populations remain well connected; however, populations from these ranges remain separated from the northern populations as seen in the California mountain kingsnake (Myers et al., 2013). Nevertheless, these geographic barriers not only restrict gene flow in the host species but can also affect gene flow in endosymbiont lineages that disperse with their hosts (Mikheyev et al., 2008).

Our ADMIXTURE analysis showed evidence of mixed ancestry in two individuals from southern and western populations, and one individual showed ancestry from all three populations (Figure 1B), implying potential gene flow among these populations. Following Pleistocene climatic cycles, species that shared glacial refugia began dispersing and expanding their ranges; this brought previously reproductively isolated populations or species into secondary contact, forming suture zones. (Hewitt, 2000; Remington, 1968; Swenson & Howard, 2004, 2005). Remington (1968) proposed three putative suture zones across California: (a) the northern California and southern Oregon, (b) the central Sierra Nevada in California and western Nevada, and (c) the southern California. Later, Swenson (2010) performed contact zone clustering analysis no evidence of contact zone in southern California. However, our observed occurrence of admixed *C. laevigatus* individuals in southern California does not overlap with Remington’s suture zones.

### Post-glacial range expansion impacts intraspecific genetic diversity in *C*. *laevigatus*

In our analysis, we observed a differential pattern of genetic variation across the *C*. *laevigatus* range. We observed reduced genetic diversity in northern populations of *C*. *laevigatus*, with a significant negative relationship between latitude and genetic diversity (Figure 2). This pattern well aligns with postglacial northward expansion. During the LGM, colder climate forced many North American taxa to contract their ranges into refugia (Batchelor et al., 2019; Boria & Blois, 2023; Kleman et al., 2010), leading to demographic changes that ultimately affected intraspecific genetic diversity (Fonseca et al., 2023; Hewitt, 2004). Genetic diversity tends to be higher in populations that persisted in refugia and maintained a stable and larger effective population size. In contrast, post-glacial range expansion towards higher latitudes often resulted in serial founder effects, smaller effective population sizes, and reduced genetic diversity in newly colonized areas (Adams & Hadly, 2013; Hewitt, 2000; Hewitt, 2004; Provan & Bennett, 2008). These processes may result in a latitudinal gradient of intraspecific genetic diversity, as observed across many North American vertebrate taxa that were co-distributed during the LGM (Adams & Hadly, 2013). For example, the California vole (*Microtus californicus*) (Conroy & Neuwald, 2008), Western Rattlesnakes (*Crotalus oreganus*) (Schmidt et al., 2020), and the kangaroo rat (genus *Dipodomys*) (Jezkova et al., 2015) all exhibited relatively high genetic diversity at lower latitudes, implying the role of postglacial range expansion on genetic diversity. Consistent with prior studies, our results also showed a similar pattern of higher genetic diversity at lower latitudes (Figure 2). Overall, we believe that southern California supported larger and stable populations of *C*. *laevigatus*, which later contributed to postglacial colonization, resulting in low genetic diversity in the direction of expansion. Lastly, our demographic analyses demonstrated that the initial split occurred during the early Pleistocene and subsequent divergence occurred during the mid Pleistocene, highlighting the role of Quaternary climatic cycles in the diversification of *C*. *laevigatus* lineages (Figure 6).

### Codiversification between *C*. *laevigatus* and *Blochmaniella*

Since *Blochmaniella* is maternally inherited and vertically transmitted, we expected their phylogeographic patterns to mirror their hosts. As expected, on a broad scale, our cophylogenetic analysis indeed revealed a largely congruent pattern between the host and endosymbiont phylogenies, supporting the codiversification hypothesis (Figure 3). Phylogenetic congruence, or codiversification, is expected between a host and its vertically transmitted endosymbiont (Wernegreen, 2002) and similar patterns have consistently been observed in insect-bacterial associations, such as plant bugs, sap-feeding bugs, mealybugs (Downie & Gullan, 2005; Fromont et al., 2016; Moran et al., 2005; Wang et al., 2018; Wernegreen, 2002), weevils (Toju et al., 2013), bees (Koch et al., 2013), and cockroaches (Garrick et al., 2017).

In contrast, incomplete lineage sorting (ILS) can arise when ancestral polymorphisms are retained across diverging populations, leading to discordance between species and gene trees (Degnan & Rosenberg, 2009; Hardig et al., 2000; Martins et al., 2009). This may be the case we observed in the northern population, particularly in those from the Sierra Nevada and Klamath Mountains, where we identified phylogenetic incongruence between the host and its endosymbiont. The observed phylogenetic incongruence between the host and its maternally inherited endosymbiont is not surprising, as similar patterns have been documented in previous studies. For example, a study by Symula et al. (2011) on the tsetse fly and its symbiont *Wigglesworthia* revealed phylogeographic congruence on a broad geographic scale; however, at smaller regional scales, discordance was likely due to incomplete lineage sorting. However, ILS alone does not fully account for the observed discordance; other factors, such as climatic shifts and dispersal differences, can independently generate discordance between host and endosymbiont phylogenies by affecting demographic processes. For instance, Garrick et al. (2017) suggested that stochastic processes, such as genetic drift or a strong bottleneck, may contribute to discordance between cockroaches and their endosymbiont, *Blattabacterium*.

Moreover, this discordance between host and endosymbiont phylogenies can be explained by climatic forcing and sex dispersal differences. As the climate grew colder during the Pleistocene, these conditions forced several species to seek refuge (Batchelor et al., 2019; Boria & Blois, 2023; Kleman et al., 2010). These glacial refugia have played a key role in lineage diversification by providing suitable habitats during the glacial maxima (Stewart et al., 2010). For instance, the Klamath Mountains and the Sierra Nevada served as glacial refugia for several taxa (Eckert et al., 2008; Goebel et al., 2009; Olson et al., 2012; Schierenbeck, 2014; Shafer et al., 2010). Having said that, the *C*. *laevigatus* populations in these regions likely survived through glacial periods, maintaining highly structured *Blochmaniella* lineages. Alternatively, differences in selective pressures acting on mito-/endosymbiont genomes versus host nuclear genomes, long-term geographic isolation, and sex-biased dispersal could also cause the phylogenetic incongruence (Funk & Omland, 2003; Symula et al., 2011; Toews & Brelsford, 2012). Specifically, female-biased dispersal may explain the pattern seen in *Blochmaniella*, given their maternal inheritance. Together, these factors may explain the discrepancy between the hosts’ and endosymbionts’ phylogenetic trees and EEMS analysis within genetic clusters.

Overall, the host and endosymbiont both followed a pattern of isolation by distance, indicating a strong positive correlation between genetic and geographic distance across all three lineages (Figure 4A-B). These results revealed that IBD was a key factor associated with the spatial patterns of intraspecific divergence among the three populations. Such patterns are commonly reported in phylogeographic studies. For example, in California, similar IBD patterns have been found in low-dispersal terrestrial taxa, like the California slender salamander (*Batrachoseps attenuatus*) (Martinez-Solano et al., 2007), the dusky-footed woodrat (*Neotoma fuscipes*) (Boria et al., 2021), the California newt (*Taricha torosa*) (Kuchta & Tan, 2006), in a mygalomorph spider (*Antrodiaetus riversi*) (Hedin et al., 2013), as well as in low-dispersal avian taxa such as the wrentit (*Chamaea fasciata*) (Burns & Barhoum, 2006) and the California thrasher (*Toxostoma redivivum*) (Sgariglia & Burns, 2003). These species, like our focal species, show continuous distributions in northern California, extending into the northern Coast Ranges and the Sierra Nevada foothills, with the Central Valley acting as a major phylogeographic barrier. Altogether, our results suggest that the genetic structure that we observe is shaped not only by physical barriers but also by limited dispersal. Furthermore, the EEMS analysis supports these findings by highlighting regions with low effective migration surface across known biogeographic barriers, such as the Central Valley and the Mojave Desert.

### Kinship

In eusocial insects such as ants, patterns of genetic relatedness are often associated with colony structure and mating strategies (Hölldobler & Wilson, 1990). Comprehending these familial relationships within and between colonies may help shed light on how dispersal dynamics and social structure operate (Crozier & Pamilo, 1996). However, such studies have received less attention in the genus *Camponotus* despite their diversity compared to the less diverse genus *Formica* (Meadows et al., 2023). Earlier studies have suggested that the genus *Camponotus* exhibits a monogynous colony structure due to independent colony formation and worker caste polymorphism (Crozier & Pamilo, 1996; Goodisman & Hahn, 2005; Hölldobler & Wilson, 1990). This is further supported by Meadows et al. (2023) reporting that within-colony worker-worker relatedness and kinship are generally high across *C. herculeanus*, *C. laevissimus,* and *C. modoc,* reflecting a high intra-colony genetic relatedness pattern.

Despite the small sample size, our relatedness analysis based on R0 and KING-robust kinship ratios showed that several foragers collected from the same oak tree were full-sisters, consistent with previous studies. For example, Meadows et al. (2023) utilized a small sample size and found full-sister relationships among intra-colony individuals. Interestingly, three individuals in our analysis—C-260, C-264b, and C-285a—did not display full-sister relationships with co-foragers, despite being collected while foraging with the other sampled individuals (i.e., C-260a, C-260b; C-264, C-264a; C-285, C-285b) (Figure 5B). This observed pattern is unsurprising, as we sampled foragers rather than nest-bound colony members. We speculate that these individuals may have belonged to different colonies or shared overlapping foraging areas; therefore, they ended up together on the same oak tree due to their foraging behavior. Such behavior has been observed in carpenter ants, where foragers often leave the nest independently in search of food (Yamamoto & Del-Claro, 2008). It is also possible that some colonies have multiply mated queens, which can result in distantly related individuals; however, we could not confirm this because of our sampling scheme. Additionally, comparisons between ants within 1 km (nearby trees) revealed that most pairwise comparisons were half-siblings (Figure 5B-D), suggesting that the queens founding these colonies are likely sisters or closely related.

## Acknowledgements

We thank Brandon Meadows and Ari Rice for their assistance during field work. This work was supported by National Science Foundation Grant 2238571 and Texas Tech University start-up funds to JDM. Mohamed Fokar at the TTU Center for Biotechnology & Genomics provided sequencing support. The TTU Center for Biotechnology & Genomics acquisition of the NovaSeq6000 was supported by NIH grant 1S10OD025115-01. We are thankful to the High-Performance Computing Center at Texas Tech University for providing computational resources.

## Data Accessibility and Benefit-Sharing

Raw sequence reads are deposited in the NCBI SRA repository with an accession under PRJNA1297169. The code used in this study can be found on GitHub: https://github.com/SwapnilBoyane/Camponotus_laevigatus_Phylogeography

## Author Contributions

J.D.M. conceived the idea for the study and acquired funding; J.D.M. and G.J.B. contributed to sample collection and performed DNA extractions. S.S.B. performed the formal analyses with partial analytical support from G.J.B. and input from J.D.M. S.S.B. wrote the original draft with feedback from J.D.M. All authors reviewed and approved the final version of the manuscript.

