## Supplemental Table 1 for "Quaternary climatic changes and biogeographic barriers drove codiversification in the obligate mutualism between *Camponotus laevigatus* and its endosymbiont *Blochmaniella*"

Supplementary Table 1. Sampling localities of individuals used in this study. Accession numbers will be based on the sampling site and not per individual.

| Colony_id | Species | Lineage | Latitude | Longitude | Accession numbers (to be populated) | Genomic coverage of *C. laevigatus* | Genomic coverage of *Blochmaniella* |
|---|---|---|---|---|---|---|---|
| C-142 | *C. laevigatus* | South | 32.77773 | -116.44528 | | 7.20 | 646.28 |
| C-145 | *C. laevigatus* | South | 32.77671 | -116.44923 | | 16.61 | 275.28 |
| C-148 | *C. laevigatus* | South | 34.38662 | -117.77512 | | 24.18 | 428.06 |
| C-150 | *C. laevigatus* | South | 34.38776 | -117.77746 | | 30.57 | 635.12 |
| C-260 | *C. laevigatus* | South | 33.65237 | -117.44998 | | 24.30 | 1194.31 |
| C-260a | *C. laevigatus* | South | 33.65237 | -117.44998 | | 23.15 | 580.51 |
| C-260b | *C. laevigatus* | South | 33.65237 | -117.44998 | | 22.92 | 1341.73 |
| C-261 | *C. laevigatus* | South | 33.65297 | -117.44847 | | 27.18 | 2453.43 |
| C-264 | *C. laevigatus* | West | 35.26102 | -120.41492 | | 20.45 | 306.72 |
| C-264a | *C. laevigatus* | West | 35.26102 | -120.41492 | | 26.22 | 854.52 |
| C-264b | *C. laevigatus* | West | 35.26102 | -120.41492 | | 36.83 | 1043.91 |
| C-265 | *C. laevigatus* | West | 35.26587 | -120.40801 | | 33.39 | 849.32 |
| C-266 | *C. laevigatus* | West | 36.23186 | -121.48379 | | 7.68 | 239.99 |
| C-267 | *C. laevigatus* | West | 36.23417 | -121.48138 | | 27.17 | 1003.26 |
| C-167 | *C. laevigatus* | North | 41.92675 | -122.83006 | | 16.23 | 304.18 |
| C-269 | *C. laevigatus* | North | 39.43986 | -122.96917 | | 20.86 | 529.40 |
| C-270 | *C. laevigatus* | North | 39.4419 | -122.97079 | | 128.94 | 608.65 |
| C-270a | *C. laevigatus* | North | 39.4419 | -122.97079 | | 26.18 | 536.98 |
| C-270b | *C. laevigatus* | North | 39.4419 | -122.97079 | | 24.56 | 776.89 |
| C-273 | *C. laevigatus* | North | 40.81862 | -122.89975 | | 24.53 | 583.55 |
| C-284 | *C. laevigatus* | North | 38.48545 | -120.26167 | | 16.42 | 434.28 |
| C-285 | *C. laevigatus* | North | 38.48646 | -120.26102 | | 29.19 | 198.43 |
| C-285a | *C. laevigatus* | North | 38.48646 | -120.26102 | | 26.31 | 717.07 |
| C-285b | *C. laevigatus* | North | 38.48646 | -120.26102 | | 32.60 | 1100.75 |
| C-287 | *C. laevigatus* | North | 37.83857 | -120.05173 | | 19.79 | 565.15 |

| | | | | | | | |
|---|---|---|---|---|---|---|---|
| C-289 | *C. laevigatus* | North | 37.83807 | -120.04684 | | 18.36 | 0.77 |
| C-291 | *C. laevigatus* | North | 37.03783 | -119.24033 | | 22.61 | 529.81 |
| C-293 | *C. laevigatus* | North | 37.03572 | -119.23914 | | 21.53 | 382.85 |
| C-294 | *C. laevigatus* | North | 35.96276 | -118.47807 | | 24.89 | 1010.81 |
| SRR19335438 | *C. modoc* | Outgroup | - | - | - | 27.11 | 272.46 |
